# The solute carrier MFSD1 decreases β1 integrin’s activation status and thus tumor metastasis

**DOI:** 10.1101/2021.09.17.460743

**Authors:** Marko Roblek, Julia Bicher, Merel van Gogh, Attila György, Rita Seeböck, Bozena Szulc, Markus Damme, Mariusz Olczak, Lubor Borsig, Daria Siekhaus

## Abstract

Solute carriers are increasingly recognized as participating in a plethora of pathologies, including cancer. We describe here the involvement of the orphan solute carrier MFSD1 in the regulation of tumor cell migration. Loss of MFSD1 enabled higher levels of metastasis in a mouse model. We identified an increased migratory potential in MFSD1^-/-^ tumor cells which was mediated by increased focal adhesion turn-over, reduced stability of mature inactive β1 integrin, and the resulting increased integrin activation index. We show that MFSD1 promoted recycling to the cell surface of endocytosed inactive β1 integrin and thereby protected β1 integrin from proteolytic degradation; this led to dampening of the integrin activation index. Furthermore, down-regulation of MFSD1 expression was observed during early steps of tumorigenesis and higher MFSD1 expression levels correlate with a better cancer patient prognosis. In sum, we describe a requirement for endolysosomal MFSD1 in efficient β1 integrin recycling to suppress tumor spread.

## 1 Introduction

Solute carriers (SLCs) represent the second most numerous class of integral membrane proteins, coded by 456 known genes [1], and are outnumbered only by G-protein coupled receptors. SLCs transport a broad spectrum of different molecules (e.g. sugars, neurotransmitters, vitamins, nucleosides, amino acids, etc.) and are therefore essential for the maintenance of homeostasis in mammalian cells, tissues and organs. About 100 SLC genes are known to cause Mendelian disorders upon mutation [2], while more are expected to have roles in multigenic diseases [1]. Interest in studying SLCs is increasing due to their roles in diverse diseases such as neurological disorders, metabolic syndrome, cardiovascular diseases, and cancer [3–6]. As of now, only 19 SLC proteins are targeted by drugs [7], and more than 30% of SLCs remain orphans, with their physiological substrates and functions unknown [1].

Previously we have described the essential function of Minerva (Mrva), the fly ortholog of the SLC Major Facilitator Superfamily Domain-containing protein 1 (MFSD1), for the invasive migration of macrophages in the developing *D. melanogaster* embryo [8]. The re-expression of mouse MFSD1 in macrophages of *mrva^-/-^* embryos rescued their invasive cell migration, indicating functional conservation from fly to mouse and a cell-autonomous capacity to control migration by MFSD1 [8]. MFSD1 was initially identified as a lysosomal membrane protein [9], and was additionally shown to be required for normal tissue physiology [10], as MFSD1^-/-^ mice develop splenomegaly and severe liver disease. MFSD1’s stability in lysosomes depends on its interaction with Glycosylated Lysosomal Membrane Protein (GLMP) [9, 11]. Even these few examples show that MFSD1 is crucial for normal physiology, however its cellular and molecular functions remain unknown.

Given MFSD1’s capacity to regulate the invasive migration of macrophages in the fly embryo, we were interested in whether MFSD1 is involved in mammalian tumor cell migration. *Drosophila* macrophages require integrin for their invasive migration [12]. Tumor cell migration depends to a huge extent on integrins, which mediate the anchorage of the cell to the ECM and relay mechanical force to the actin cytoskeleton [13]. The deregulation of integrins by diverse mechanisms contributes to a pro-tumorigenic and pro-metastatic phenotype [13], by affecting expression levels, endocytosis and recycling dynamics [14, 15], and glycosylation status [16, 17].

Here we show that MFSD1 in tumor cells restrains metastasis, decreases the rate of migration, and lowers the resistance to mechanical stress and starvation induced apoptosis by reducing the activation index of β1 integrin through the recycling pathway.

## 2 Materials and Methods

### Cell culture

Murine colon carcinoma cell line MC-38 [18], melanoma cell line B16-BL6 [19], and breast cancer cell line 4T1 (ATCC, #CRL-2539) were cultured in DMEM (ThermoFisher Scientific, 31966-021) supplemented with 10% FCS (Sigma, #9665) and 1x NEAA (ThermoFisher Scientific, #11140-050) and cultivated in incubators set at 37°C and 5% CO_2_.

MFSD1^kd^ tumor cells were generated by MISSION lentiviral transduction particles expressing short hairpin RNA (shRNA) from pLKO.1 vector targeting the coding sequence of MFSD1 (Sigma, TRC clone ID TRCN0000338002 and TRCN0000337937) or control shRNA (Sigma, #SHC216V). Infected cells were selected by Puromycin (2 μg/ml) for five days.

MFSD1^-/-^ and cognate WT cells were generated from an MC-38 single cell clone progenitor (gift from H. Clausen lab) and transiently transfected with Lipofectamin 3000 according to manufacturer’s instructions with gRNA targeting MFSD1 (MMPD0000065783, gRNA sequence: 5’-GGCGGTGTTCCCGTTCATC-3’) or GLMP (MMPD0000057180, gRNA sequence: 5’-ACTTGGCCAAGGAGTACGG-3’) in line with GFP from the (p04) U6-gRNA:CMV-eCas9-2a-tGFP plasmid (Sigma). One day later, GFP^+^ cells have been single cell sorted into 96-well plates by a BD Aria III sorter (BD). Single cell clones have been expanded and evaluated for efficient knock-out by Western blotting. Five single cell clones for MFSD1^-/-^ and WT, respectively, were combined for subsequent experiments. MC-38 Cosmc^-/-^ cells were a gift from H. Clausen.

### Recombinant proteins

The cloning of MFSD1-eGFP was described previously [8]. GLMP-HA was PCR amplified from pcDNA3.1-GLMP-HA [11] with forward primer 5’-GGGGACAAGTTTGTACAAAAAAGCAGGCTTAATGTTTCGCTGTTGG-3’ and reverse primer 5’-GGGGACCACTTTGTACAAGAAAGCTGGGTATTAAGCGTAGTCTGGGAC-3’. The PCR product was cloned using Gateway BP Clonase II Enzyme mix and Gateway LR Clonase II Enyzme Mix (ThermoFisher Scientific) via donor vector pDonR211 into the final Doxycycline inducible expression vector pInducer20 [20] according to manufacturer’s instructions. QSOX1-StrepTagII was PCR amplified from pCMV3-C-Myc-QSOX1 (Sinobiological, #MG53456-CM) with forward primer 5’-GGGGACAAGTTTGTACAAAAAAGCAGGCTTAAAGCTTGGTACCATG-3’ and reverse primer 5’-GGGGACCACTTTGTACAAGAAAGCTGGGTATTACTTTTCGAACTGCGGGTGGCTCCAAGAGCCTCCACCCCC-3′. The PCR product was processed as was the GLMP-HA construct. All constructs were separately packed into lentiviral particles using pdelta8.9 (Addgene, #2221) and pCMV-VSV-G (Addgene, #8454) packaging plamid transfected Lenti-X 293T cells (TaKaRa, #632180) using Lipofectamine 3000 (ThermoFisher Scientific). Crude lentiviral supernatant was used for infection of MC-38 cells.

### Mice

All animal experiments were done according to the guidelines of the Swiss Animal Protection Law and approved by the Veterinary Office of the Kanton Zürich. C57BL/6J mice were purchased from The Jackson Laboratory and maintained in individually ventilated cages at 21°C, 55% humidity, with a photoperiod of 12 hours light and 12 hours dark with standard diet and water *ad libidum*.

### Western blot

Cells were lysed with lysis buffer (25 mM Tris, 150 mM NaCl, 1 mM EDTA, 1% Triton X-100, supplemented with Halt Protease and Phosphatase Inhibitor Cocktail (ThermoFisher Scientific, #78440) for 20 min on ice and cleared by centrifugation for 10 min at 20’000 x g. Cell lysates (10 μg) were separated on 4-15% SDS-PAGE gradient gels (Bio-Rad) and blotted on Protran 0.45 nitrocellulose membranes (GE Healthcare). Membranes were blocked with 1x Pierce Clear Milk Blocking Buffer (ThermoFisher Scientific, #37587) for 1 h at RT. Primary antibodies were diluted in 1x blocking buffer and incubated overnight at 4°C and included: anti-MFSD1 [10], anti-β tubulin (Cell Signaling Technology, #2146S), anti-GLMP [10], anti-β1 integrin (ThermoFisher Scientific, #PA5-78028), anti-GAPDH (Abcam, #ab181603), anti-T antigen (clone 3C9, gift from H. Clausen, [21]), anti-QSOX1 (gift from D. Fass, [22]), anti-LAMP1 (clone 1D4B, BioLegend, #121601), anti-cleaved caspase 3 (Cell Signaling Technology, #9661S), Streptavidin-HRP (ThermoFisher Scientific, #S911). Subsequently, membranes were washed 3x with TBS-T and incubated with either Goat Anti-Mouse IgG (H+L)-HRP Conjugate (Bio-Rad, #1721011) or Goat Anti-Rabbit IgG (H+L)-HRP Conjugate (Bio-Rad, #1706515) secondary antibody. After 3x washing with TBS-T, the membrane was incubated with SuperSignal West Femto Maximum Sensitivity Substrate (ThermoFisher Scientific, #34096) and the chemoluminescence signals were detected with the ChemiDoc MP Gel Imaging System (Bio-Rad). Densitometric analysis of Western blot bands was performed with ImageJ.

### Immunofluorescence

The expression of MFSD1-eGFP was induced by 100 ng/μl Doxycycline for 24 h prior to cell fixation. Cells were seeded on Nunc Lab-Tek Chamber Slides (ThermoFisher Scientific, #154534) and fixed with 4% FA/PBS (ThermoFisher Scientific, #28906) for 10 min at RT. Samples were washed 3x with PBS and blocked/permeabilized with 1%BSA/0.1%Triton X-100 in PBS for 1 h at RT. Samples were then stained with antibodies for 2 h at RT, which included: anti-GFP (clone 5G4, from E. Ogris), anti-HA-tag (clone 3F10, Sigma, #11867432001), anti-giantin (BioLegend, #19243), LysoTracker Red DND-99 (ThermoFisher Scientific, #L7528), anti-Rab5A (Cell Signaling Technologies, #46449), anti-Rab7 (Cell Signaling Technologies, #9367), anti-Rab11 (Cell Signaling Technologies, #5589), anti-mTOR (Cell Signaling Technologies, #2983S), anti-paxillin (Abcam, #ab32084). Samples were washed 3x with PBS and incubated with secondary antibodies, which included: anti-mouse AlexaFluor488 (ThermoFisher Scientific, #A11001), anti-rabbit AlexaFluor555 (ThermoFisher Scientific, #A31572), anti-rabbit AlexaFluor488 (ThermoFisher Scientific, #A11008), anti-rabbit AlexaFluor633 (ThermoFisher Scientific, #A21070), anti-rat AlexaFluor546 (ThermoFisher Scientific, #A11081). Samples were washed and counter stained with DAPI for 10 min and subsequently mounted in ProLong Diamond Antifade Mountant (ThermoFisher Scientific, #P36970). Pictures were taken on a Zeiss LSM880 inv. Fast Airyscan or Zeiss LSM800 inv. Airyscan confocal microscopes with a Plan-Apochromat 40x/NA 1.3 OIL, or Plan-Apochromat 40x/1.2 Water objective, respectively. Colocalization (Pearson’s R value), fluorescence intensity, and particle size was analyzed by ImageJ.

### Flow cytometry

24 h before staining, cells were incubated with or without 10 μg/ml 9EG7 antibody (BD Biosciences, #553715), or 20 μg/ml Primaquine (Sigma, #160393). Cells were stained in 100 μl FACS buffer (2% FCS/10mM EDTA in PBS) with following antibodies and lectins: PNA-FITC, WGA-FITC, and LPA-FITC were from EY laboratories (FITC-labeled lectin kit #2), anti-β1 integrin (clone HMb1-1, BioLegend, #102202), anti-β1 integrin-APC (clone HMb1-1, BioLegend, #102215), anti-active β1 integrin (clone 9EG7, BD Biosciences, #553715), 7-AAD (ThermoFisher Scientific, #A1310), AnnexinV-PE (BioLegend, #640908). Cells were washed once with 1 ml buffer and, if applicable, stained again in 100 μl buffer with the anti-rat AlexaFluor488 (ThermoFisher Scientific, #A11006) secondary antibody, followed by one wash with 1 ml buffer. Cells were resuspended in 300 μl buffer and data were acquired on a BD FACS Canto II Analyzer (BD Biosciences) and analyzed by FlowJo software (TreeStar Inc.).

### Wound closing assay

60’000 cells were seeded per chamber on Culture-Insert 2 Well in μ-Dish 35 mm (ibidi GmbH) 24 h before treatment with the cytostatic Mitomycin C (10 μg/ml, Sigma, #M4287) for 3.5 h in normal medium, followed by removal of the insert, and one wash step with cell culture medium. For start of the migration fresh cell culture medium was added with or without 10 μg/ml 9EG7 antibody (BD Biosciences, #553715). The progress of migration was recorded every minute with temperature and CO2 controlled Leica DM IL LED microscopes with Leica HI PLAN I 10x/0.22 air objectives. Videos were analyzed with Wound Healing FastTrack AI Image Analysis (ibidi GesmbH), or respective images at indicated time points are shown. The wound size at the start of migration was equalized to the smallest wound size at the particular experiment to correct for different wound sizes.

### Experimental metastasis

300’000 MC-38 or 150’000 B16-BL6 cells in 100 μl HBSS (Sigma, #H9394) were injected into the tail vein of C57BL/6J mice. Four or two weeks after intravenous injection of tumor cells, respectively, lungs were perfused with PBS and the number of macroscopic metastatic foci was counted.

### Matrigel invasion assay

100’000 MC-38 cells were seeded in 500 μl DMEM + 1% FCS atop of Corning BioCoat Matrigel Invasion Chambers with 8 μm PET Membrane (Corning, #354480) and chemotaxis was induced by presence of DMEM + 10% FCS in the bottom chamber. Invasive migration was stopped after 24 h and cells reaching the bottom side of the PET Membrane were fixed with methanol and stained with Giemsa’s solution. PET membranes were cut out and mounted in ProLong Diamond Antifade Mountant (ThermoFisher Scientific, #P36970) on objective slides. Per membrane 3-4 pictures at a random location were taken for determination of the efficiency of invasive migration with a Leica DMI 6000B Microscope with a Leica N PLAN 10x/0.25 air objective.

### Proliferation ratio determination

MC-38 WT-eGFP and MC-38 MFSD1^-/-^-mCherry cells were seeded together in 6-well plates and their relative ratio was determined for the zero hour time-point. This ratio was observed also after 24, 48, 72, and 96 h after seeding. No change of ratio indicates equal proliferation.

### Spreading assay

The experiment was done as described in [23]. Briefly, cells were starved of serum for 1 h prior to two minutes incubation with Trypsin-EDTA (ThermoFisher Scientific, #25300-054), quenched with serum-free medium and centrifuged for 5 min at 300 x g. Cells were resuspended in a cell density of 200’000 cells/ml in serum-free medium and incubated in the conical tube in the incubator. Meanwhile, 50 μl of serum-free medium was added to wells of a 96-well plate, which have or have not been previously coated with 10 μg/ml fibronectin (Sigma, #F2006) for 30 min, and equilibrated in the incubator. Then 50 μl of the cell suspension was added and incubated for 20 and 40 min followed by direct addition of 20 μl 25% Glutaraldehyde (Sigma, #G5882). Cells were fixed for 30 min at RT and then the fixed cells were washed and stored in PBS. Three random pictures at each replicate (individual 96-well) were taken by a Leica DM IL LED microscope with Leica HI PLAN I 10x/0.22 air objective. Spread cells were manually counted.

### CORA analysis

To analyze *O*-glycans, procedure reported by Kudelka et al. [24] was adopted, as reported recently [25, 26]. In brief, the cells were incubated in medium containing 5% serum with 50 μM peracetylated *O*-glycan precursor (Ac_3_GalNAcBn) and grown for 72 h. Then, the conditioned media were collected, centrifuged (1000 × g, 5 min), and the supernatants were subjected to glycan extraction procedure. Next, isolated O-glycans were permethylated and subsequently analyzed using MALDI-TOF mass spectrometry.

### N-glycan analysis

Cells from 10 cm plates were lysed using the CelLytic M reagent (Sigma-Aldrich), and the protein concentration in obtained lysates was adjusted to 2 mg/mL. Cellular proteins were precipitated by the addition of one volume of iced 100% acetone and incubated at −20 °C overnight. Samples were centrifuged (14000g, 15 min, RT) and the pellet containing precipitated proteins was air-dried. Precipitated proteins were resuspended in 150 μL of Glycoprotein Denaturing Buffer (New England Biolabs) and incubated at 100 °C for 10 min. Afterwards, the following PNGase F reaction mixture was assembled: cellular protein sample (100 μL), 10% NP-40 (20 μL, NEB), G7 Reaction buffer (20 μL, NEB), deionized water (60 μL) and PNGase F (4.5 μL, 150 U). The reaction mixture was incubated at 37 °C for 4 h, afterwards another 3 μL of PNGase F were added and incubated for a total of 24 h at 37 °C. Purification of N-glycans from the reaction mixture was done on the Supelclean™ ENVI-Carb™ SPE graphitized carbon tubes (Sigma-Aldrich). Prior to purification, the columns were treated sequentially with 1 M NaOH (3 mL), H_2_O (2 x 3 mL), 30% acetic acid (3 mL), and H_2_O (2×3 mL). Afterwards, 3 mL of 50% acetonitrile with 0.05% TFA was added, followed by 6 mL of 3% acetonitrile with 0.05% TFA. Glycan samples were centrifuged and the supernatant was applied to the columns. The columns were then washed first with water (3 mL), then with 3% acetonitrile and 0.05% TFA solution. Glycans were eluted with 50% acetonitrile and 0.05% TFA solution (4 x 0.5 ml). Obtained samples were lyophilized overnight in a vacuum concentrator.

Released *N*-glycans were fluorescently labeled on the non-reducing end with 2-aminobenzamide (2-AB) and separated on GlycoSepN column (Prozyme) connected to HPLC system, as previously described [27].

### MALDI-TOF analysis of N- and O-glycans

MALDI-TOF-MS experiments were performed on an Axima-Performance TOF spectrometer (Shimadzu Biotech), equipped with a nitrogen laser (337 nm). The pulsed extraction ion source accelerated the ions to a kinetic energy of 20 keV. Data were obtained in a positive-ion linear mode. The calibration of the linear mode analysis was done using polyethylene glycol in mass range up to 5000 Da. The accuracy of the product ion calibration is approximately 1.5 Da. The mass calibration was conducted based on the average masses. The samples were dissolved in 20% acetonitrile in water. As a matrix, 2,5-dihydroxybenzoic acid (20 mg/mL) dissolved in 20 mM sodium acetate in 20% acetonitrile in water was used. The sample and matrix were combined at a 1:1 ratio. The resulting solution (1 μl) was spotted on a 384-well MALDI-TOF plate, followed by evaporation of the solvent at ambient temperature without any assistance. Each mass spectrum was accumulated from at least 200 laser shots and processed by Biotech Launchpad ver. 2.9.1 program (Shimadzu).

### Detachment experiment

Cells were grown on uncoated or fibronectin coated μ-Slide 8-well chambers (ibidi, #80826) to 80-90% confluency, washed with PBS and detached with 10 mM EDTA in PBS, 230 μM RGDS peptide (Tocris, #3498) in PBS, or 0.05% Trypsin/EDTA (ThermoFisher Scientific, #25300-054). Detachment was observed under microscope and within few minutes (2-4 min) slides were rocked and detaching solution was sucked off and immediately remaining bound cells were fixed with 4% FA/PBS for 15 min. Fixed cells were washed and stained with crystal violet for 30 min at RT, washed again and stored in PBS. 3-4 pictures at random locations of each chamber was taken with a Leica DM IL LED microscope with Leica HI PLAN I 10x/0.22 air objective. Area covered by still adherent cells was determined with ImageJ.

### QSOX1 activity assay

QSOX1-STII was purified from the supernatant of MC-38 WT or MFSD1^-/-^ cells and 0.5 μg was used in a total reaction volume of 150 μl in 96-well plates. Final reaction mixture contained 1mM Homovanilic acid (Sigma, #1252), 1.4μM horseradish peroxidase (Sigma, #P6782), 133μM DTT, 50mm K_2_HPO_4_ (pH7.5), 1mM EDTA. Purified QSOX1-STII was added to the wells just prior start of measurements. Dimerization of Homovanilic acid was recorded every 30 seconds for 20 minutes with a BioTek Synergy H1 platereader (BioTek) spectrophotometer with an Ex = 320 nm and Em = 420 nm.

### RNA isolation and qPCR

RNA was isolated with the RNeasy Mini Kit (Qiagen, #74104) and cDNA produced with the Omniscript RT Kit (Qiagen, #205111) with oligo dT primers. qPCR reactions were assembled according to the Takyon No ROX SYBR 2x MasterMix blue dTTP (Eurogentec, #UF-NSMT-B0701) qPCR kit. qPCR was performed on a LightCycler 480 (Roche) and data were analyzed with the Lightcycler 480 software. 2^−ΔΔct^ (compared to WT) are displayed. The following primers were used: GAPDH-fw: CCCAGCAAGGACACTGAGCAA; GAPDH-rev: GTGGGTGCAGCGAACTTTATTGATG; ITGb1-fw: GAAAGCAGGGAGAAGTTGGC; ITGb1-rev: TGATGTCGGGACCAGTAGGA.

### Immunoprecipitation of β1 integrin incl. glycosidase experiment

β1 integrin was immunoprecipitated with anti-β1 integrin (clone HMb1-1, BioLegend, #102202), or anti-active β1 integrin (clone 9EG7, BD Biosciences, #553715) from cell lysates for 30 min on ice, followed by pull-down with Dynabeads Protein G (ThermoFisher Scientific, #10003D) for 1 h on ice. Beads were washed 3x with PBS and either boiled in 1x Laemmli Sample Buffer (Bio-Rad, #1610747), or treated with EndoH (New England BioLabs, #P0702S) or PNGaseF (New England BioLabs, #P0704S) according to manufacturer’s instructions.

Cell surface β1 integrin was immunoprecipitated with incubation of intact cells with the aforementioned antibodies in FACS buffer for 30 min on ice, washed once with PBS, lysed with 1% Triton X-100 in PBS. Cell lysate was cleared by centrifugation and the protocol continued as described in the previous paragraph.

### Starvation experiment

Cells were grown till they reached 80-90% confluency. Cell culture medium was removed, cells were washed twice with PBS and were incubated for 48 h in DMEM (ThermoFisher Scientific, 31966-021) with or without 10 μg/ml 9EG7 antibody (BD Biosciences, #553715) or 0.16 μM Bafilomycin A1 (Sigma, #SML1661). Cells were lysed with lysis buffer and proceeded for analysis by Western blotting.

### Mechanical stress resistance experiment

Cells were grown on 6-well plates until 80-90% confluency in presence or absence of 10 μg/ml 9EG7 antibody (BD Biosciences, #553715) for 24 h and scraped off manually with a rubber Corning Cell Lifter (Corning, #3008). Cells were stained with 7-AAD (ThermoFisher Scientific, #A1310) and AnnexinV-PE (BioLegend, #640908) and data were acquired by flow cytometry with a BD FACS Canto II Analyzer (BD Biosciences) and analyzed by FlowJo software (TreeStar Inc.).

### Cell surface biotinylation experiment

Cells were grown on 6-well plates until 80-90% confluency and cell surface proteins were biotinylated with EZ-Link NHS-PEG4-Biotin (ThermoFisher Scientific, #21330) at 0.5 mg/ml in HBSS (ThermoFisher Scientific, #14170112) for 45 min at 4°C. Biotin was washed off with PBS and cells were either lysed (zero hour time point) or incubated with normal cell culture medium for 12 h and then lysed (12 h time point) with cell lysis buffer. Cell lysates were cleared by centrifugation and β1 integrin was immunoprecipitated with anti-β1 integrin (clone HMb1-1, BioLegend, #102202) and Dynabeads Protein G (ThermoFisher Scientific, #10003D) for 15 min at RT. Beads were washed twice with PBS and then boiled in 1x Laemmli Sample Buffer (Bio-Rad, #1610747). Samples were subjected to Western blot analysis.

### Degradation of β1 integrin

Cells were grown on 6-well plates until 80-90% confluency. Cells were then treated with 5 μg/ml Brefeldin A (Sigma, #B7651) for 12 h in normal cell culture medium and then lysed. Cell lysates were analyzed by Western blot.

### Statistical analysis

Statistical analysis was performed with GraphPad Prism software (version 9.2.0). Data are presented as mean ± SEM and were analyzed depending on the number of groups by 2way ANOVA with the Šidak’s multiple comparison test or with the Mann-Whitney test.

## 3 Results

### MFSD1 slows tumor cell migration and metastasis initiation

We aimed to study the effect of MFSD1 on tumor cell migration. We prepared a murine colon carcinoma cell line MC-38 MFSD1^kd^ stably expressing a shRNA which reduces MFSD1 expression by 75% compared to the levels in corresponding control cells expressing a nt-shRNA (Figure 1A). We achieved similar levels of shRNA knock-down of MFSD1 in two other murine tumor cell lines, 4T1 breast cancer and B16-BL6 melanoma (Suppl. Figure 1A). MC-38 MFSD1^kd^ cells migrated faster in a wound closing assay compared to cognate control cells (Figure 1B–C). This increased migration was also observed upon MFSD1 knockdown in 4T1 and B16-BL6 tumor cell lines (Suppl Figure 1B), supporting a role for MFSD1 in tumor cell migration across tumor cell types. We also observed increased invasive migration by MC-38 MFSD1^kd^ cells in a Matrigel invasion assay (Suppl. Figure 1C). Next we sought to expand our *in vitro* tumor migration assays into *in vivo* metastasis. We detected increased number of lung metastases in mice *i.v.* injected with MFSD1^kd^ tumor cells compared to the control for both MC-38 (Figure 1D) and B16-BL6 (Suppl. Figure 1D) cells. To eliminate the residual MFSD1 expression seen in the MFSD1^kd^ condition we generated MC-38 MFSD1^-/-^ cells by CRISPR-Cas9 mediated gene editing (Suppl. Fig 1E). MFSD1^-/-^ cells closed the wound faster in the wound closing assays when compared to respective WT cells (Figure 1E–F) confirming the MFSD1^kd^ data. The proliferation of cells was unaffected by the presence of MFSD1 (Suppl. Figure 1H), indicating that the difference in migration and metastatic foci formation is independent of an effect on proliferation. Thus these experiments support the conclusion that in multiple types of murine cancer cell lines MFSD1 suppresses general and invasive migration *in vitro* and metastasis *in vivo*.

**Figure 1.**
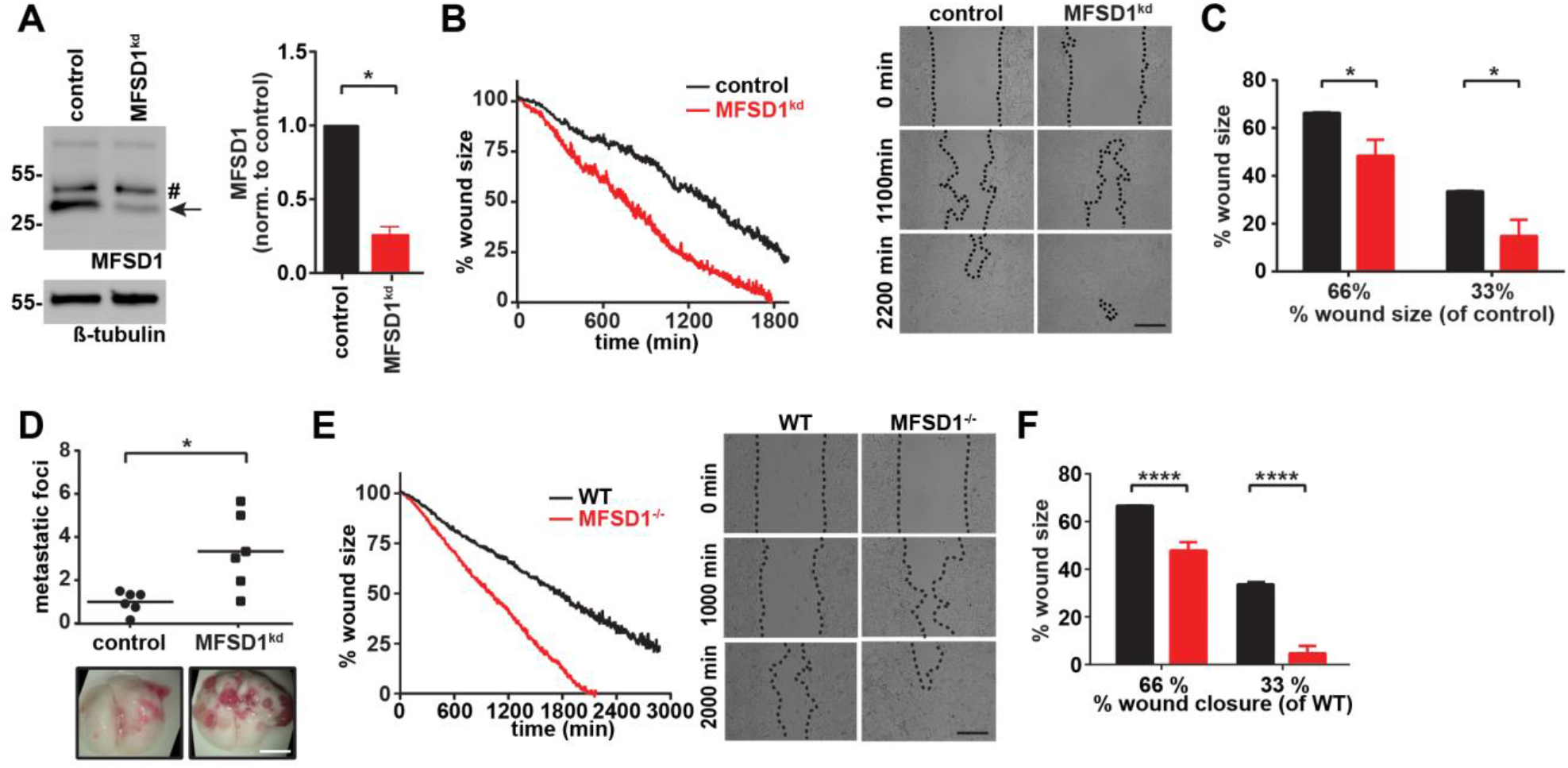
MFSD1 suppresses cell migration. **A)** Western blot of MC-38 cells expressing shRNA non-target (control) or shRNA MFSD1 (MFSD1^kd^) (left panel). Quantification of MFSD1 protein levels by densitometry (right panel) (n=4). Arrow indicates MFSD1 band, # indicates unspecific band. **B)** Wound closing assay of MC-38 control and MFSD1^kd^ cells. Graph depicting the continuous shrinking of the wound (left panel) with pictures at defined time-points (right panel). Data from one representative experiment are shown. Bar = 0.2 mm. **C)** Analysis of the wound closing assay of MC-38 control and MFSD1^kd^ cells (n=4). The % wound size remaining for MFSD1^kd^ cells when control cells have moved to shrink the wound to either 66% or 33% of the original size is depicted. **D)** Experimental metastasis with MC-38 control and MFSD1^kd^ cells. The relative number (normalized to the mean seen in the control) of macroscopic metastatic foci per lung is shown (n=6). Representative images of lungs are shown below. Scale bar = 0.5 cm. **E)** Wound closing assay of MC-38 WT and MFSD1^-/-^ cells. Graph depicting the continuous shrinking of the wound (left panel) with pictures at defined time-points (right panel). Data from one representative experiment are shown. Bar = 0.2 mm. **F)** Analysis of wound closing assay of MC-38 WT and MFSD1^-/-^ cells (n=5). The % wound size remaining for MFSD1^-/-^ cells when control cells have moved to shrink the wound to either 66% or 33% of the original size is depicted. * = p<0.05; **** = p<0.0001.

### MFSD1 requires the accessory subunit GLMP for its migratory function

Previously GLMP has been identified as a crucial interaction partner of MFSD1 [10], thus we additionally prepared MC-38 GLMP^-/-^ cells with the CRISPR-Cas9 method. Western blotting with respective antibodies confirmed the elimination of the endogenous protein in the MFSD1^-/-^ and GLMP^-/-^ lines and we could confirm that the stability of the two proteins is mutually dependent (Suppl. Figure 1E). GLMP^-/-^ cells also migrated and closed the wound faster than corresponding WT cells (Suppl. Figure F–G), phenocopying the MFSD1^-/-^ cells.

### MFSD1 localizes to the Golgi and to the endolysosomal system

To gain insight into the possible functions of MFSD1, we examined its sub-cellular location in tumor cells. Previously, we had shown that it localizes predominantly to the Golgi apparatus in tumor cell lines, but is found in both the Golgi apparatus and the endolysosomal system in fly macrophages [8]. To enable a deeper analysis of its localization we used Doxycycline-inducible expression of MFSD1-eGFP (Suppl. Figure 2A), as constitutive expression of MFSD1 was toxic to cells (data not shown). In tumor cells the strongest MFSD1 staining co-localized with its accessory subunit GLMP in the Golgi apparatus (Figure 2A). However, a careful scrutiny of MFSD1 stainings revealed an additional weaker vesicle-like staining pattern. This MFSD1-eGFP^+^ pattern co-localized with acidic compartments stained in live conditions by the LysoTracker dye, indicating MFSD1’s presence within the endolysosomal system (Figure 2A–B). To further delimit MFSD1 to specific endosomal vesicles we stained cells with a panel of vesicle specific antibodies, including the early endosomal marker Rab5, late endosomal marker Rab7, recycling endosomal marker Rab11, and lysosomal marker mTOR (Suppl. Figure 2B). MFSD1-eGFP co-localized with the late endosomal marker Rab7 and the lysosomal marker mTOR, and partially with the recycling endosomal marker Rab11 (Suppl. Figure 2B). To assess if MFSD1’s localization pattern is conserved across different cell types we examined MFSD1-eGFP expressed in NIH-3T3 fibroblasts. Interestingly, here we found MFSD1 positioned exclusively in the endolysosomal compartment (Suppl. Figure 2C), localized with the same sub-cellular markers as in tumor cells (co-localization with Rab7 and mTOR, and partial co-localization with Rab11) (Figure 2C and Suppl. Figure 2C–D). The same additional weaker vesicle-like staining pattern seen with MFSD1-eGFP was also observed for GLMP, indicating their co-localization in the Golgi apparatus as well as in the endolysosomal compartment (Figure 2A and Suppl. Figure 2E).

**Figure 2.**
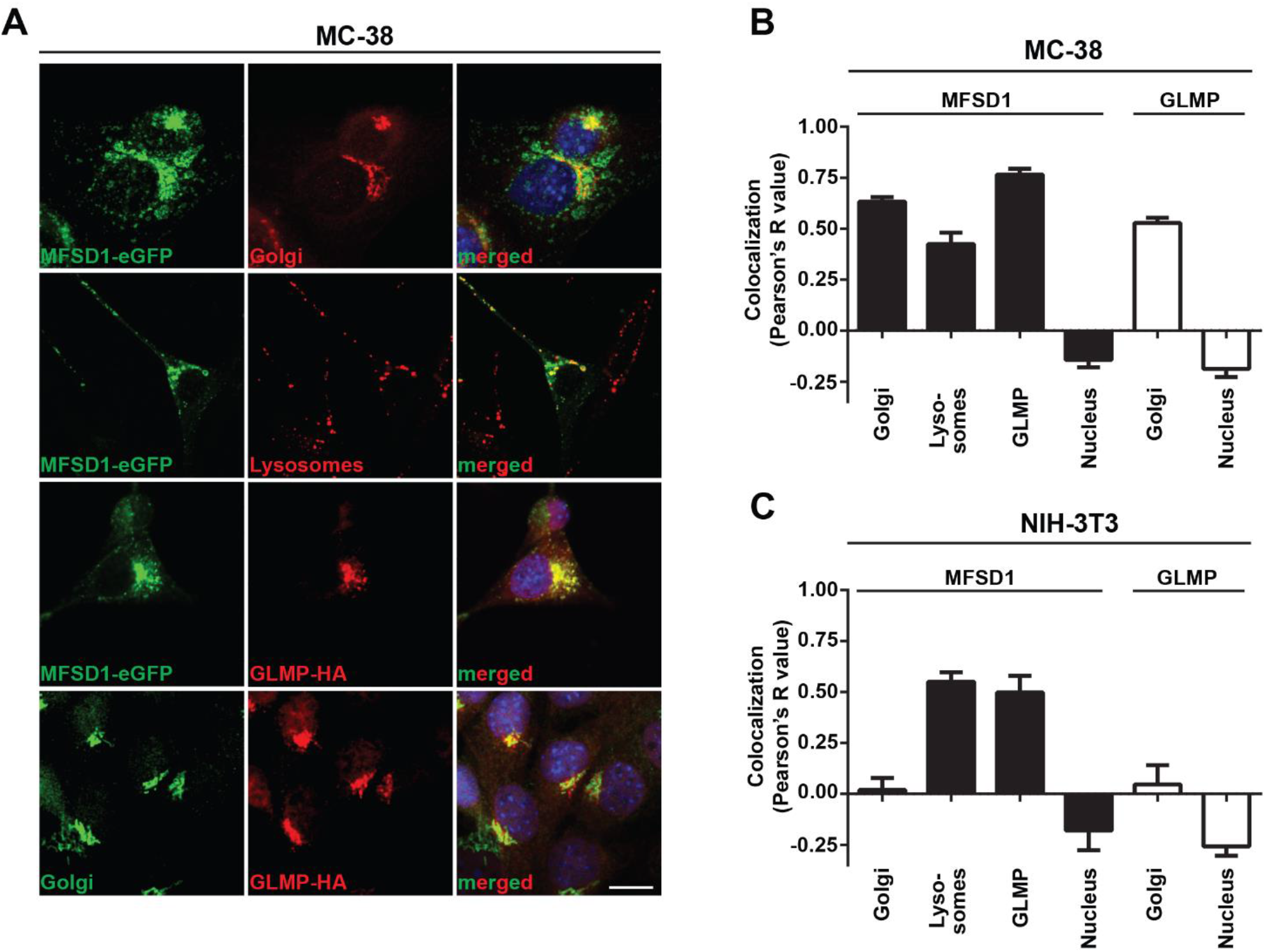
MFSD1-eGFP localizes to the Golgi apparatus and endo-lysosomal system in tumor cells. **A)** Immunofluorescence pictures of MC-38 MFSD1-eGFP cells with the indicated markers. Bar = 10 μm. **B)** Co-localization analysis of MFSD1-eGFP and GLMP-HA with indicated sub-cellular markers in MC-38 tumor cells (n≥13). **C)** Co-localization analysis of MFSD1-eGFP and GLMP-HA with indicated sub-cellular markers in NIH-3T3 fibroblasts (n≥4).

We hypothesize that the Golgi localization of MFSD1-eGFP in tumor cells might be due to tumor cell-specific retention in the Golgi apparatus. MFSD1’s presence in tumors in the Golgi apparatus and the endolysosomal system indicate two different axes through which it could affect biological processes, which we addressed in our subsequent experiments.

### MFSD1 does not influence glycosylation

Cell surface glycans, added to proteins in the Golgi apparatus, play important roles during cell migration and metastatic spread [28, 29]. Previously we have described an effect of Minerva, the fly orthologue of MFSD1, on the O-glycome of *D. melanogaster* embryos [8]. We tested the functionality of the O-glycosylation machinery after the initial GalNAc addition in our tumor cells by the Cellular O-Glycome Reporter/Amplification (CORA) approach [24]. This assay uses a modified GalNAc-α-Benzyl which cannot be covalently attached to proteins, but which is secreted to the medium upon extension with other sugars by the action of glycosyl-transferases. The overall secreted O-glycome of MC-38 WT and MFSD1^-/-^ cells consisted predominantly of sialyl- and disialyl-T antigen, which were present in the same amounts in both cell types (Suppl. Figure 3A), indicating that the enzymes required for O-glycan generation subsequent to initial GalNAc addition are present and functional in both cell types. In addition, we tested the presence of the O-glycan T antigen on the enzyme QSOX1, whose level we showed previously to be affected by the fly’s MFSD1 homologue [8]. Secreted QSOX1-StrepTagII from MC-38 WT, MFSD1^-/-^ and Cosmc^-/-^ cells (which lack the functional T-synthase required for the generation of T antigen) was treated by sialidase to remove the terminal sialic acid, which allowed the detection of the T antigen by a specific antibody. While Cosmc^-/-^ cells did not have any T antigen, the levels of T antigen were indistinguishable between WT and MFSD1^-/-^ cells (Suppl. Figure 3B). In addition, the enzymatic activity of QSOX1-StrepTagII purified from MC-38 WT and MFSD1^-/-^ cells was indistinguishable (Suppl. Figure 3C). By liquid chromatography, we tested for differences in the N-glycan pool and found essentially identical retention profiles of N-glycans isolated from WT and MFSD1^-/-^ cells (Suppl. Figure 3D). As an example, the N-glycosylation of the heavily N-glycosylated protein LAMP1 was not altered in MFSD1^-/-^ compared to WT cells (Suppl. Figure 3E). We tested lectin binding to the cell surface of MC-38 WT and MFSD1^-/-^ cells and could detect sialic acid (LPA), GlcNAc (WGA), and Gal (PNA) sugar-containing glycans on the cell surface, with the same abundance in both MC-38 genotypes (Suppl. Figure 3F–G). From these experiments we conclude that most likely neither O- nor N-glycosylation are directly affected by MFSD1 in tumor cells.

### MFSD1 has minor effects on lysosomal function

We continued with a basic characterization of the endolysosomal system in MC-38 WT and MFSD1^-/-^ cells stained by LysoTracker. The size of the lysosomes in MFSD1^-/-^ was reduced by 9%, while the activity of Cathepsin B, a protease used to measure lysosomal functionality, was increased by 26 %, compared to WT lysosomes (Suppl. Figure 3H). The total number of lysosomes per cell and the pH in the endolysosomal system was not affected by the lack of MFSD1 (Suppl. Figure 3H).

### MFSD1 affects β1-integrin and adhesion dynamics of tumor cells

In cell culture, we observed faster detachment of MC-38 MFSD1^kd^ and MFSD1^-/-^ compared to WT MC-38 cells (Suppl. Figure 3I–J) indicating weaker adhesion to the cell culture plates. We also observed faster spreading of MFSD1^-/-^ cells on uncoated and fibronectin-coated plates over time when compared to WT MC-38 cells (Figure 3A and Suppl. Figure 3K), suggesting MFSD1^-/-^ cells have a faster turn-over of binding sites, a condition which has previously been shown to promote metastasis [30]. Therefore, we analyzed focal adhesions in MC-38 WT and MFSD1^-/-^ cells by immunofluorescent staining of paxillin. Both paxillin fluorescent intensity (FI) and the average size of paxillin stained focal adhesions were reduced in MFSD1^-/-^ compared to WT cells, while the total number of focal adhesions per cell remained the same (Figure 3C–D). Focal adhesions link the actin cytoskeleton to the extracellular matrix via integrins. When we performed a Western blot of MC-38 WT and MFSD1^-/-^ cells we observed a double band of β1-integrin, which changed in its relative intensity (Figure 3E). When we quantified the β1-integrin levels by densitometry we detected that the overall amount of β1-integrin was decreased in MFSD1^-/-^ when compared to WT cells (Figure 3F), while its mRNA levels was not affected (Suppl. Figure 3L). The level of the immature N-glycosylated lower β1-integrin band was unaffected, but the level of the mature N-glycosylated upper β1-integrin band was decreased in MFSD1^-/-^ when compared to WT MC-38 cells (Figure 3F). Thus we observed that MFSD1 stabilizes focal adhesions and the maturely N-glycosylated β1-integrin protein in MC-38 tumor cells.

**Figure 3.**
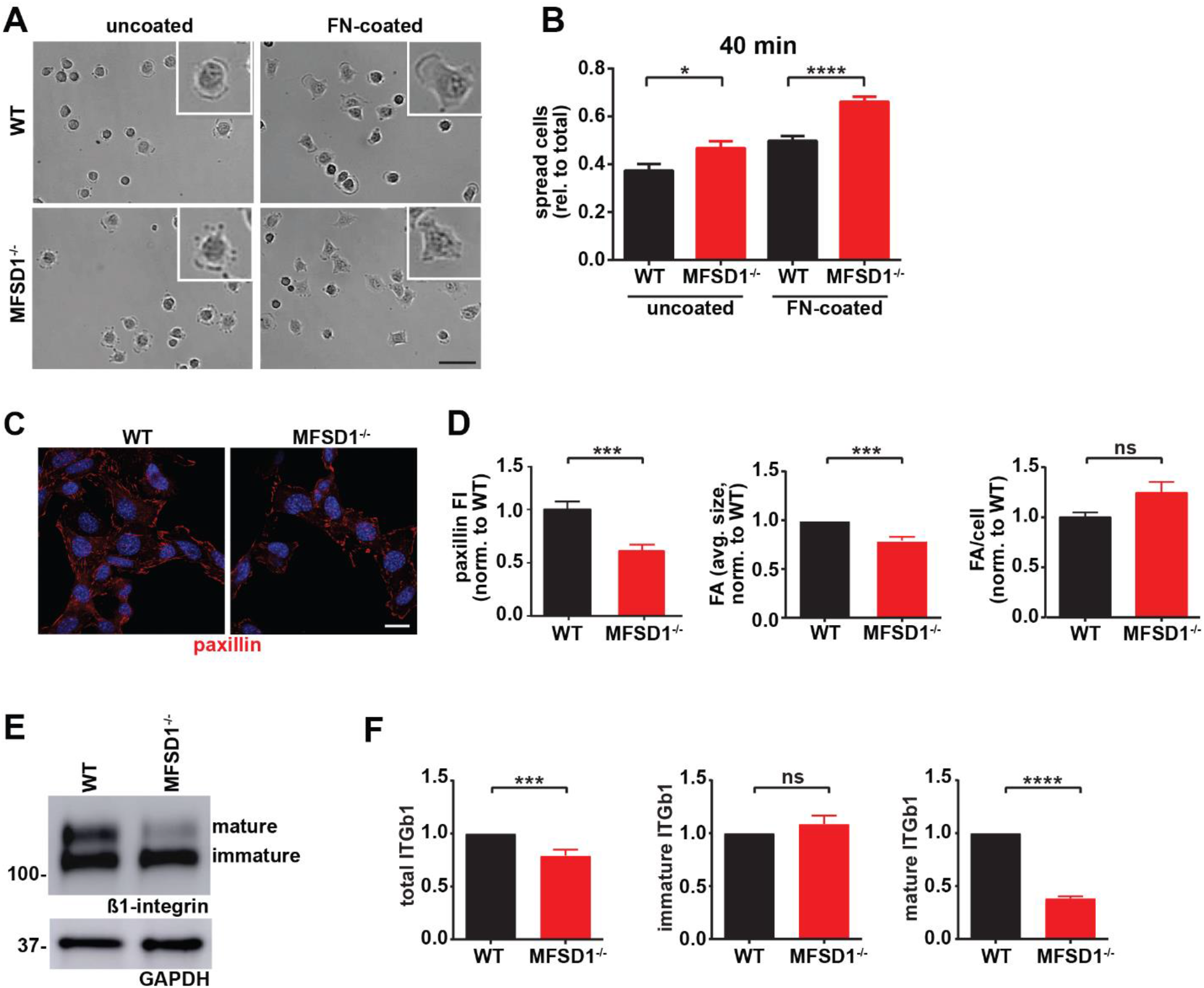
MFSD1 controls cell adhesion dynamics. **A)** Spreading assay of MC-38 WT and MFSD1^-/-^ on uncoated and FN-coated plates. Representative images after 40 min of spreading are shown. Magnified spread cell in the top right corner of each view field is shown. Bar = 50 μm. **B)** Quantification of spreading assay at 40 min time point. Experiment was performed three times in duplicate or triplicate and three view fields were analyzed per condition (n≥21 view fields). **C)** Immunofluorescence of focal adhesions with paxillin staining in MC-38 WT and MFSD1^-/-^ cells. Bar = 10 μm. **D)** Analysis of paxillin immunofluorescence staining. **E)** Western blot of MC-38 WT and MFSD1^-/-^ cells for β1-integrin. GAPDH serves as a loading control. **F)** Densitometric analysis of β1-integrin detected by Western blots (n=10). * = p<0.05; *** = p<0.001; **** = p<0.0001; ns = not significant.

### MFSD1 enhances the stability of maturely N-glycosylated, inactive β1-integrin

We wished to further characterize the activation state of the different β1-integrin forms we had observed in Western blots. We, therefore, performed immunoprecipitation experiments on lysates from MC-38 WT and MFSD1^-/-^ cells prepared from total cells or just the cell surface. While the conformation insensitive β1-integrin antibody clone HMb1-1 detected both forms of β1-integrin, the active conformation-specific β1-integrin antibody clone 9EG7 specifically immunoprecipitated only the lower β1-integrin band from both samples (Figure 4A). We sought to test if the different sizes of the β1-integrin bands are due to differential N-glycan modification. Therefore, we treated lysates from MC-38 WT and MFSD1^-/-^ cells from the total cell and the intact cell surface with PNGaseF to remove all N-glycans. Immunoprecipitation with an antibody that recognizes all forms of β1-integrin shows that PNGaseF treatment de-glycosylated both bands and yields a single band of non-glycosylated β1-integrin at 88 kDa in all samples (Suppl. Figure 4A). Furthermore, the lower β1-integrin band was indeed sensitive to EndoH treatment, being reduced to the size of the form lacking all glycosylation, indicating that the lower β1-integrin band corresponds to an immaturely N-glycosylated form. This result highlights that immaturely glycosylated β1-integrin makes it to the cell surface, and is predominantly in its active conformation. Summarizing, our data show that MFSD1 stabilizes maturely N-glycosylated inactive β1-integrin.

**Figure 4.**
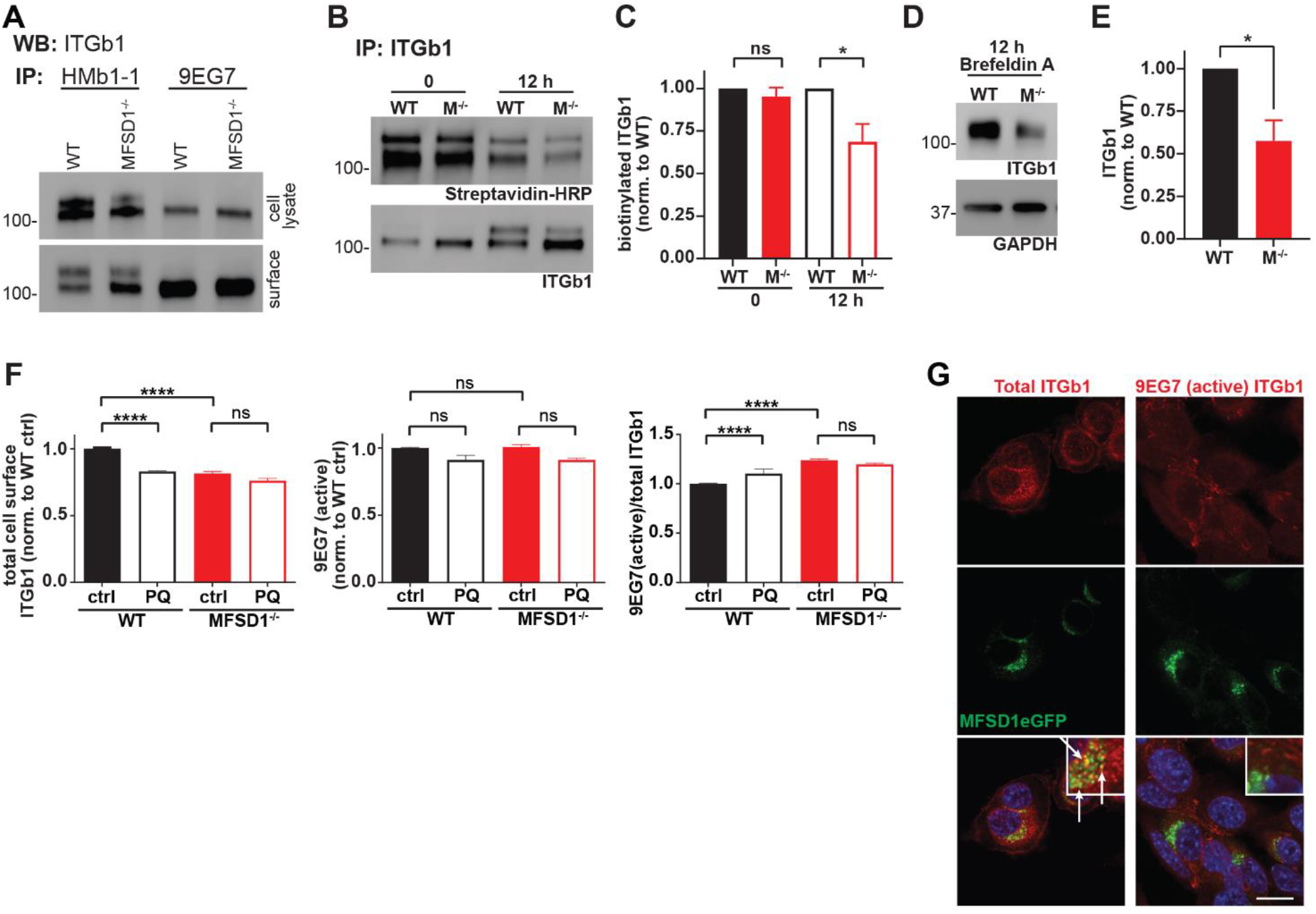
MFSD1 promotes recycling and stabilization of inactive β1-integrin. **A)** Western blot of β1-integrin immunoprecipitated by HMb1-1 or 9EG7 antibodies from total cell lysates or intact cell surface (n=2). **B)** Western blot of cell surface biotinylated and immunoprecipitated β1-integrin for biotinylation (Streptavidin-HRP) at 0 and 12 h time point post-treatment (biotinylation). ITGb1 blot displays the input of β1-integrin (n=4). **C)** Densitometric quantification of biotinylated β1-integrin (n=4). **D)** Western blot of β1-integrin upon Brefeldin A treatment for 12 h (rate of β1-integrin degradation experiment). GAPDH serves as a loading control. Representative experiment is shown. **E)** Densitometric quantification of ITGb1 from Brefeldin A treated cells for 12 h (n=4). **F)** Flow cytometric analysis of MC-38 WT and MFSD1^-/-^ cells treated with recycling inhibitor Primaquine (PQ) or control (n=7). **G)** Co-immunofluorescence of MC-38 MFSD1-eGFP cells with an antibody detecting either total (Itgb1) (n=14) or active conformation (9EG7) (n=6) β1-integrin, respectively. Bar = 10 μm.* = p<0.05; **** = p<0.0001; ns = not significant.

### MFSD1 promotes recycling and thus stabilization of inactive β1-integrin

We tested for an effect of MFSD1 on the degradation of β1-integrin in MC-38 cells. We conducted surface biotinylation and twelve hours later immunoprecipitated β1-integrin and analyzed the level of biotinylated β1-integrin that remained via Streptavidin-HRP. While the input (time point 0 hours) of biotinylated proteins was roughly the same, 12 hours later MFSD1^-/-^ cells had significantly less biotinylated β1-integrin left when compared to MC-38 WT control cells (Figure 4B–C). In addition, we blocked *de novo* feeding of proteins into the secretory pathway by treatment with Brefeldin A. Twelve hours after treatment the level of β1-integrin, which at that time was purely in the mature N-glycosylated form (Suppl. Figure 4B), was reduced to half in MC-38 MFSD1^-/-^ compared to WT cells (Figure 4D-E). Thus MFSD1 stabilizes mature β1-integrin.

To test if the recycling pathway is involved in β1-integrin stabilization, we treated MC-38 WT and MFSD1^-/-^ cells with Primaquine, an established recycling inhibitor, and analyzed cell surface β1-integrin levels by flow cytometry. Primaquine treatment reduced cell surface levels of β1-integrin in MC-38 WT cells to the levels seen in MFSD1^-/-^ and increased the β1-integrin activation index (ratio of active/total β1-integrin). Strikingly in MFSD1^-/-^ cells themselves, Primaquine treatment did not further reduce the cell surface levels of β1-integrin or affect the activation index (Figure 4F). Consistent with this finding, we observed co-localization of total but not active β1-integrin with MFSD1 (Figure 4G), arguing that MFSD1 primarily localizes with the inactive form. This data supports the conclusion that MFSD1 enhances the recycling of β1-integrin to the cell surface, particularly the mature N-glycosylated inactive form, and protects it from lysosomal degradation.

### MFSD1 decreases the β1-integrin activation index, suppressing tumor cell migration

We wished to assess integrin activity at the cell membrane to try to connect MFSD1’s capacity to increase integrin maturation and focal adhesion size to its role in suppressing cell migration. To specifically analyze the cell surface fraction of β1-integrin, we conducted flow cytometry. We stained MC-38 cells with two antibodies detecting either total β1-integrin (clone HMb1-1) or β1-integrin in its active conformation (clone 9EG7). We observed a reduced amount of total β1-integrin on the cell surface of MC-38 MFSD1^-/-^ cells when compared to WT cells, while the amount of active β1-integrin remained the same (Figure 5A–B). This relative change leads to an increased β1-integrin activation index in MFSD1^-/-^ tumor cells, a state which has been associated with pro-metastatic features of tumor cells [14]. Thus this data supports the conclusion that MFSD1 normally reduces the β1-integrin activation index at the cell surface. Furthermore, in combination with the data from Figure 4, our data supports the conclusion that the stabilization of inactive maturely N-glycosylated β1-integrin by MFSD1 contributes to the decrease in β1-integrin’s activation index, suppressing the migration of wild-type tumor cells.

**Figure 5.**
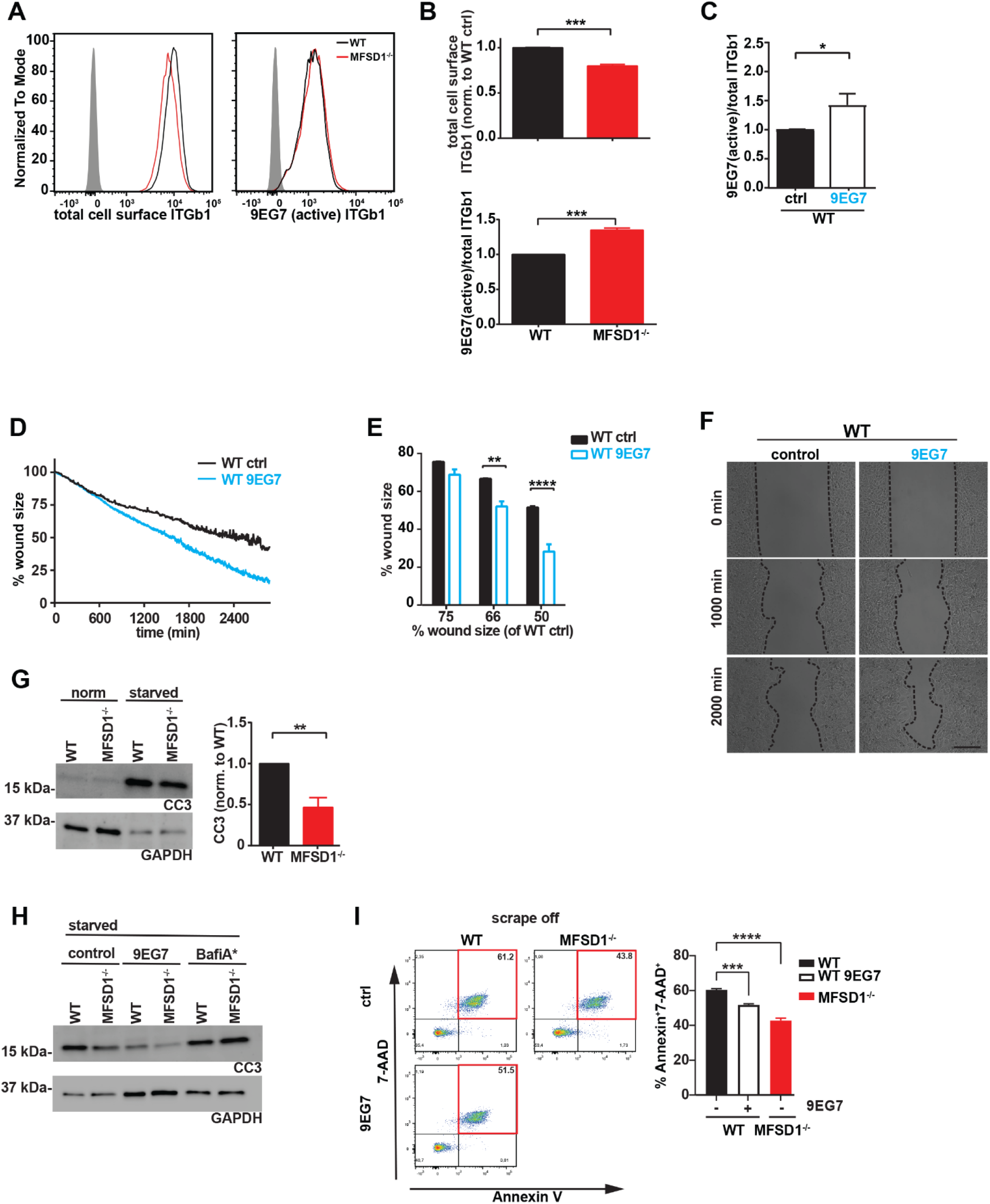
MFSD1 decreases β1-integrin activation index suppressing tumor cell migration. **A)** Representative flow cytometry analysis of MC-38 WT (black) and MFSD1^-/-^ (red) cells stained with indicated antibodies. Gray filled curve represents the isotype control staining. **B)** Analysis of flow cytometry staining (n=7). β1-integrin activation index of cells treated with active conformation specific β1-integrin antibody clone 9EG7 (n=4). **D)** Representative graph of a wound closing assay with MC-38 WT cells treated with 9EG7 antibody (dotted lines) or control (solid lines). **E)** Analysis of wound closing assay of MC-38 WT cells treated with 9EG7 antibody (empty bars) or control (filled bars) (n=3). The % wound size left of WT 9EG7 when WT cells have moved to shrink the wound to either 75%, 66%, or 50% of the original size is depicted. **F)** Representative images at indicated time points of one wound closing assay experiment. Bar = 200 μm. **G)** Western blot of cells in normal cell culture medium and in the starvation medium for cleaved-caspase 3 (CC3). GAPDH serves as a loading control (n=6). **H)** Western blot of cells previously starved from nutrients for two days, control or treated with 9EG7 antibody or Bafilomycin A (n=2). **I)** Flow cytometric analysis of dead cells (AnnexinV+7-AAD+) upon scraping (=mechanical stress) from cell culture plates. Cells have been treated with 9EG7 antibody or left untreated (control) (n≥4). * = p<0.05; ** = p<0.01; *** = p<0.001; **** = p<0.0001.

We wished to directly test the connection between β1-integrin activity and migration. We therefore incubated WT cells with the active conformation-specific β1-integrin antibody clone 9EG7. Upon 9EG7 treatment we observed an increased β1-integrin activation index as measured by flow cytometry (Figure 5C), without affecting the total amount of cell surface β1-integrin (Suppl. Figure 5A). Consistent with what we observed for MFSD1^-/-^ cells these WT 9EG7-treated cells displayed faster migration in the wound-closing assay (Figure 5D–F). This data argues that MFSD1’s suppression of the relative amount of active β1-integrin at the cell surface aids in repressing faster migration of tumor cells.

### MFSD1’s suppression of the β1-integrin activation index increases tumor cell sensitivity to nutrient starvation and mechanical stress

β1-integrin signaling positively regulates among other biological pathways the survival of tumor cells under stressed conditions, including shear stress in vascular circulation and the adaptation to changing metabolic demands [13, 14, 31, 32]. To test if the increase in the β1-integrin activation index observed in MFSD1^-/-^ cells affect their survival, we starved cells of nutrients for two days and we assessed apoptotic induction by detecting cleaved-caspase 3 (CC3) via Western blotting. As expected, MFSD1^-/-^ cells displayed less apoptotic induction compared to WT cells (Figure 5G). Interestingly, blocking lysosomal function by incubating cells with Bafilomycin A diminished the protective effect provided to the cells by the loss of MFSD1^-/-^ (Figure 5H). The starvation induced apoptosis of WT cells was rescued by incubating cells with the 9EG7 antibody to increase the β1-integrin activation index, supporting the conclusion that the effect observed in MFSD1 is due to an effect on β1-integrin. Tumor cells are exposed to shear stress in the vasculature during metastatic spread. To mimic mechanical stress, we scraped off MC-38 WT and MFSD1^-/-^ cells with a rubber cell scraper from cell culture plates and analyzed cells with apoptosis markers via flow cytometry. We observed that WT cells are more sensitive to mechanical stress-induced cell death (7-AAD^+^AnnexinV^+^ cells) than MFSD1^-/-^ cells (Figure 5I). WT cells increased their resistance to mechanical stress upon treatment with the 9EG7 antibody to increase their β1-integrin activation (Figure 5I). These results strongly suggest that MFSD1’s capacity to decrease the β1-integrin activation index in MC-38 cells is a contributing factor to its ability to suppress metastasis, by depressing tumor cell migration and increasing tumor cells’ propensity to die upon starvation or mechanical stress.

### Reduced MFSD1 expression happens early during tumorigenesis

Since we observed that MFSD1 suppresses colon cancer cell migration, we first searched for a linkage of MFSD1 expression levels and colon adenocarcinoma patients’ prognosis, but found no correlation (data not shown). Therefore, we analyzed the expression levels of MFSD1 during tumor formation. We compared publically available gene expression data from publications comparing gene expression in healthy normal mucosa and colorectal adenoma tissue, or primary colorectal tumor tissue, and found that in each of the four GEO DataSets examined [33–36], the expression levels of MFSD1 were reduced in tumor tissue when compared to healthy mucosa (Figure 6A). Thus, early reduction in MFSD1 expression might be beneficial for tumor cells to acquire hallmarks of cancer.

**Figure 6.**
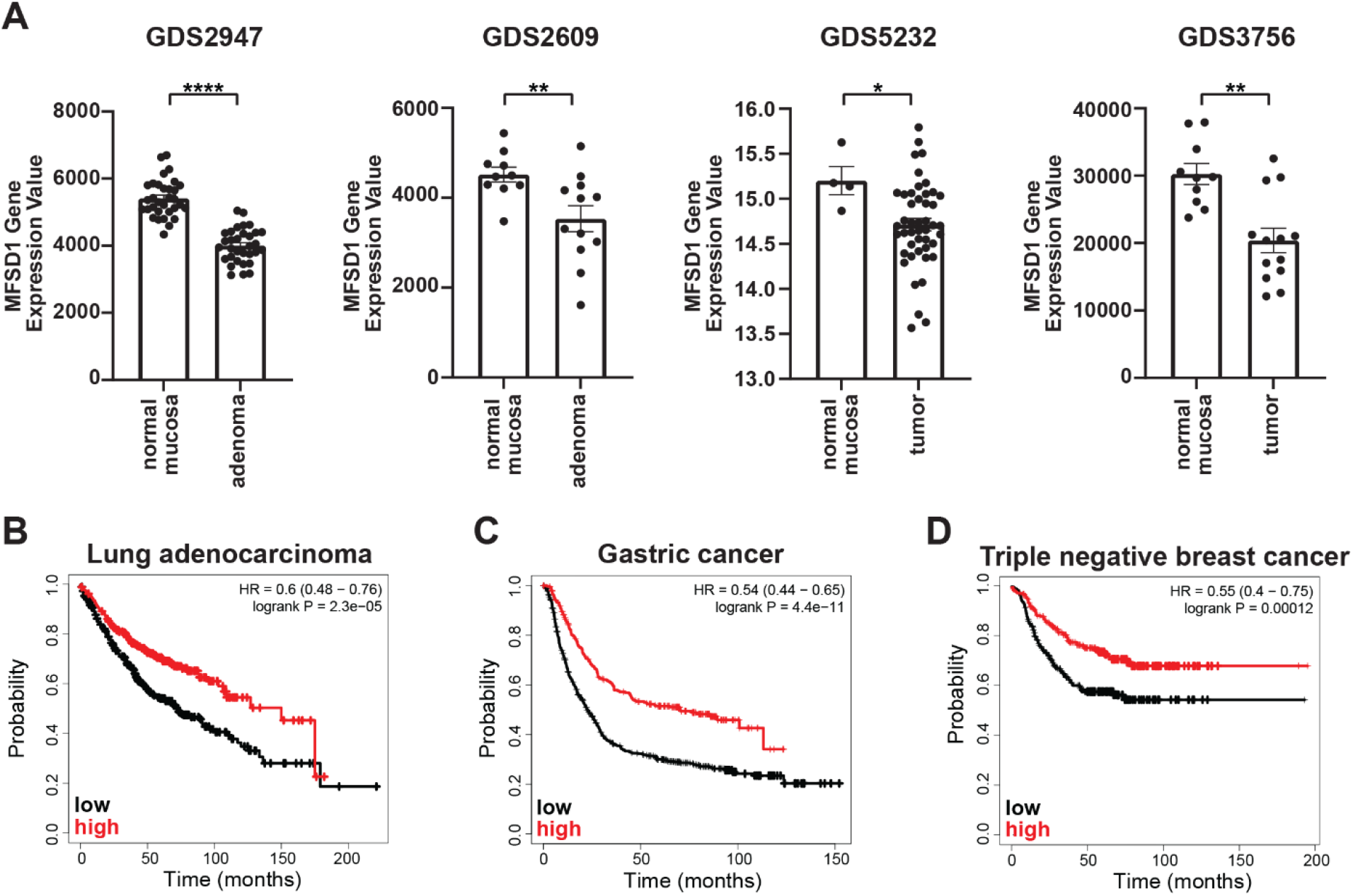
MFSD1 expression levels are reduced in colorectal adenomas and inversely correlate with patient survival in lung-, gastric-, and triple negative breast- cancer. **A)** MFSD1 gene expression values were obtained from the indicated GEO DataSets. **B-D)** KM Plotter graph of respective patient prognosis. Low or high expression levels of MFSD1 are indicated by a black or red line, respectively. * = p<0.05; ** = p<0.01; **** = p<0.0001.

### Reduced MFSD1 expression is associated with poor prognosis of selected tumor patients

We also screened data available on the Kaplan-Meier Plotter page for association of MFSD1 expression (mRNA gene chip) and the prognosis of lung-, breast-, and gastric- tumor patients [37]. Lung adenocarcinoma patients with high MFSD1 expression had a median overall survival of 150 months, compared to 72.3 months in the low expression cohort (patient group was split at the median) (Figure 6B), while lung squamous cell carcinoma patients’ prognoses were unaffected by MFSD1 expression (data not shown). The same was true for gastric cancer patients, where the median survival in the high MFDS1 expression cohort was 70.2 months and 22.5 months in the low expression cohort (Figure 6C). Refractory free survival of triple-negative breast cancer patients was significantly reduced in patients with low MFSD1 expression (Figure 6D) with a survival of the low expression cohort of 19.81 months compared to 55.2 months in the high expression cohort. These data clearly establish that MFSD1 expression correlates with patient outcome for certain tumor types.

## 4 DISCUSSION

Despite their crucial role in regulating the distribution of small molecules across membranes, the function of about one-third of SLCs remains unknown [1], including MFSD1. Here we identify a cell biological function regulated by MFSD1, namely the recycling of β1 integrin from the endolysosomal system. We show that this capacity underlies MFSD1’s regulation of migration.

MFSD1 restrained migration in the three tumor cell lines we tested. MFSD1^-/-^ tumor cells displayed smaller individual focal adhesions and increased spreading when plated on control and FN-precoated cell culture plates, as compared to WT cells. Smaller focal adhesions support faster migration [38] with a biphasic dependence of cell migration on focal adhesion size [39]. We also observed that MFSD1^-/-^ cells detached faster from cell-culture plates than WT cells, consistent with higher turn-over of focal adhesions. Increased focal adhesion turnover and focal adhesion disassembly can lead to faster *in vitro* 2D migration [40, 41] and increased incidence of *in vivo* metastasis [30, 42]. Furthermore, focal adhesion disassembly was shown to be increased in invasive versus non-invasive breast cancer cells [30]. Our findings support the conclusion that MFSD1 decreases tumor migratory speed by increasing focal adhesion size and stability.

Integrin is a key component of focal adhesions. The measure of the amount of active β1 integrin in comparison to the total β1 integrin is deemed the integrin activation index. The link between a higher integrin activity index and tumor progression or cell migration is well established [13, 14, 43–48]. For example, metastatic breast cancer and melanoma have increased activated β1 integrin compared to the respective primary tumor [44]. Furthermore, constitutively active β1 integrin L358A has an increased activation index when compared to WT β1 integrin expression and leads to increased liver metastasis of B16F10 melanoma [44]. Thus we hypothesize that MFSD1 serves as a suppressor of metastasis through its restraint of integrin activity. As we observed for MFSD1^-/-^ tumor cells, in HEK293 or MDA-MB-231 cells mutated for Shank 1 or 3, a higher β1 integrin activation index was associated with reduced adhesion area, increased cell spreading and faster migration [49]. Thus we hypothesize that MFSD1 increases focal adhesion size and stability and reduces migration speed by limiting the integrin activation index.

Cell surface integrins have a half-life of 12-24 hours and the majority of endocytosed integrin is recycled back to the cell surface [15], with active and inactive integrins using different routes and kinetics [50]. Recycling via SNX17 has been shown to be crucial for the protection of β1 integrin from lysosomal degradation [51, 52]. Differential recycling kinetics and thus protection from lysosomal degradation of the inactive or active form could lead to changes in integrin’s activation index. Our data is consistent with MFSD1’s stabilization of the inactive form of β1 integrin through stimulation of its recycling back to the cell surface, lowering the activation index. The converse action, recycling and stabilization of active β1 integrin, can be enhanced by CLIC3 or GGA2; fittingly both of these proteins stimulate the opposite phenotype for migration, namely efficient cell migration [53, 54]. We find MFSD1 localized to the endolysosomal system, just as we have observed previously for its fly ortholog, Mrva, in macrophages [8]. However, MFSD1 had a different effect on cell migration in the fly and the mammalian tumor cell system. This could be due to different effects of changes in the integrin activation on the two cell types [55, 56]. We hypothesize that MFSD1 enhances β1 integrin’s recycling and that a loss of MFSD1 leads to a changed metabolite milieu in the endolysosomal system affecting the proper function of proteins involved in the recycling of β1 integrin. Enhanced recycling of inactive β1 integrin, stimulated by MFSD1, would save β1 integrin from degradation, resulting in a reduced integrin activation index. Supporting this hypothesis is also the fact that MFSD1 co-localizes partially with recycling Rab11^+^ endosomes and total β1 integrin, and fails to co-localize with active β1 integrin.

Integrin activity is normally controlled by N-glycosylation. Removing the N-glycan site at N343 of β1 integrin greatly enhanced the fibronectin binding of the mutant α5β1 integrin compared to the control, indicating that this N-glycan site sterically impairs the active conformation or hinders ligand binding [57]. Mature glycosylated β1 integrin could be induced to have an active conformation in the presence of activating fibronectin ligand [58], indicating that N-glycosylation represses intrinsic β1 integrin activation, but allows outside-in activation. We and others have identified that immaturely N-glycosylated β1 integrin is constitutively in its active conformation [58]. We report here the presence of this form of β1 integrin on the cell surface of the murine colon carcinoma cell line MC-38 through the use of the active conformation-specific 9EG7 integrin antibody [59]. The same antibody was used to detect this immaturely glycosylated β1 integrin on the cell surface of the astrocytoma cell line A172, functionally bound to fibronectin [60]. This suggests that certain tumor cells manage to circumvent glycosylation control on β1 integrin by transporting the immaturely glycosylated form to the cell surface. Our work argues that one of the ways in which the cell suppresses tumor formation enhanced by this routing of immaturely glycosylated integrin to the cell surface is by employing MFSD1 to enhance the recycling of maturely glycosylated β1 integrin back to the cell surface to dilute out the activated forms.

MFSD1 expression levels correlate with the survival of patients with lung, breast and gastric tumors and down-regulation of MFSD1 expression happens during early events of colon tumorigenesis. Thus drugs capable of enhancing MFSD1 activity or its expression could potentially aid in inhibiting the progression of certain tumors. In future studies we aim to unravel the molecular function of MFSD1, aiding the development of drugs potentially useful for therapeutic intervention.

## 6 Author Contributions

MR and DS contributed to conception and design of the study, analyzed and interpreted data, and wrote the manuscript. MR and JB performed all but the specifically mentioned experiments. RS analyzed patient data. BS and MO performed the CORA-analysis and the N-glycan profiling. MG and LB performed in vivo metastasis experiments. MD contributed essential material. AG cloned recombinant expression constructs. All authors approved the submission of the manuscript.

## 8 Acknowledgments

We thank M Sixt, A Leithner, and J Alanko for helpful advice and the BioImaging Facility at IST Austria for technical support and assistance. We thank the Siekhaus lab for careful review of the manuscript and their input. This work was funded by the NO Forschungs- und Bildungsges.m.b.H. (LS16-021) to MR.

---

**Supplementary Figure 1.**
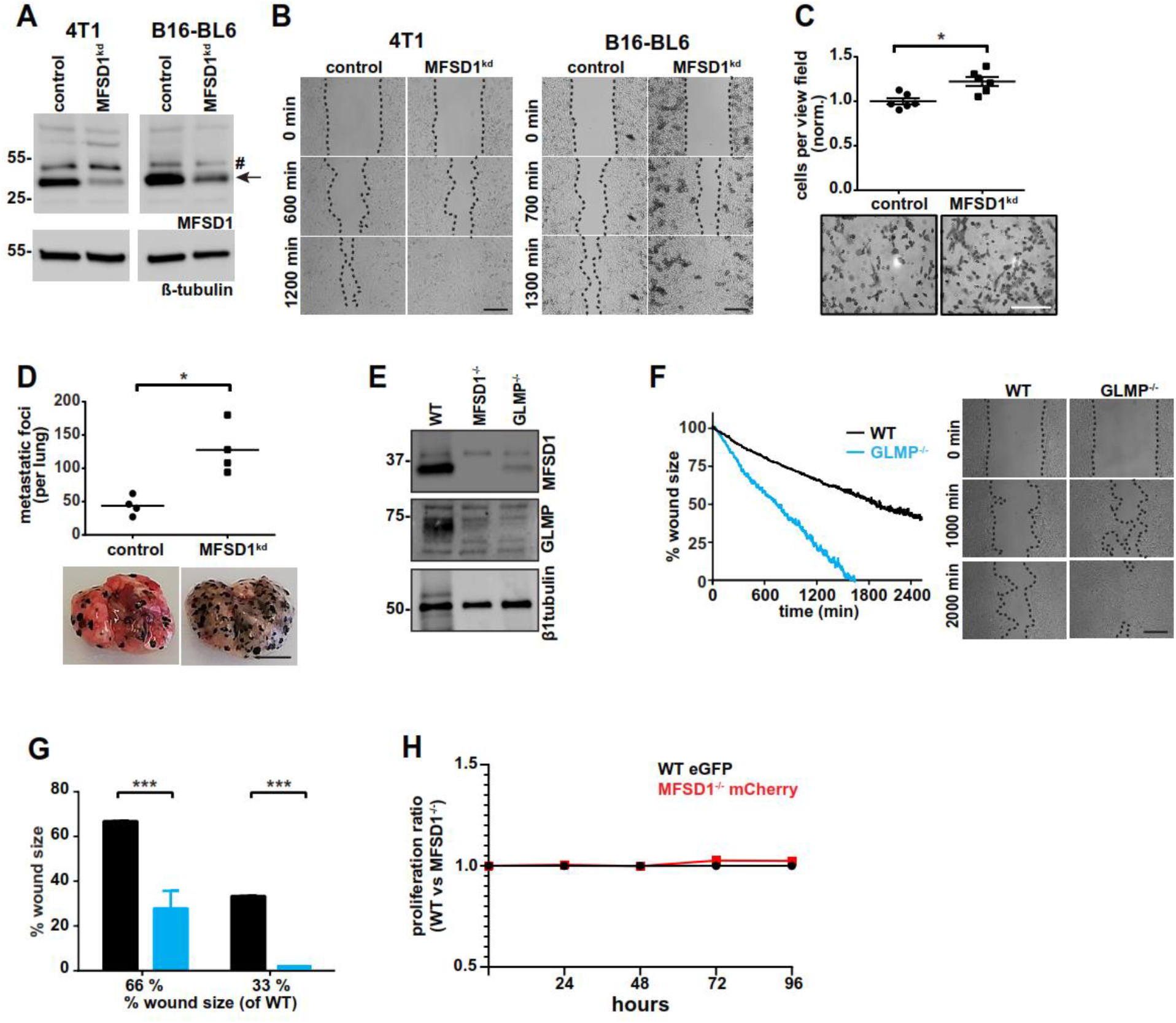
MFSD1 suppresses cell migration. **A)** Western blot of 4T1 and B16-BL6 cells expressing non-target shRNA (control) or MFSD1 shRNA (MFSD1kd). Arrow indicates MFSD1 band, # indicates unspecific band. **B)** Wound closing assay of 4T1 (left panel) and B16-BL6 (right panel) control and MFSD1^kd^ cells. Data from one representative experiment are shown (n=2). Bar = 0.2 mm. **C)** Matrigel invasion assay of MC-38 control and MFSD1^kd^ cells, with one respective view field, is shown (n=6). **D)** Experimental metastasis with B16-BL6 control and MFSD1^kd^ cells. The absolute number of macroscopic metastatic foci per lung, with representative images of lungs, is shown (n=4). Bar = 0.5 cm. **E)** Western blot of MC-38 WT, MFSD1^-/-^, and GLMP^-/-^ cells. **F)** Wound closing assay of MC-38 WT and GLMP^-/-^ cells. Graph depicting the continuous shrinking of the wound (left panel) with pictures at defined time-points (right panel). Data from one representative experiment are shown. Bar = 0.2 mm. **G)** Analysis of wound closing assay of MC-38 WT and GLMP^-/-^ cells (n=3). The % wound size left of GLMP^-/-^ cells when WT cells have moved to shrink the wound to either 66% or 33% of the original size is depicted. **H)** Proliferation index of MC-38 WT and MFSD1^-/-^ cells analyzed by flow cytometry. In this particular experiment, WT cells expressed eGFP and MFSD1^-/-^ cells expressed mCherry (n=3). Similar results were obtained with WT mCherry and MFSD1^-/-^ eGFP cells (data not shown). * = p<0.05; *** = p<0.001.

**Supplementary Figure 2.**
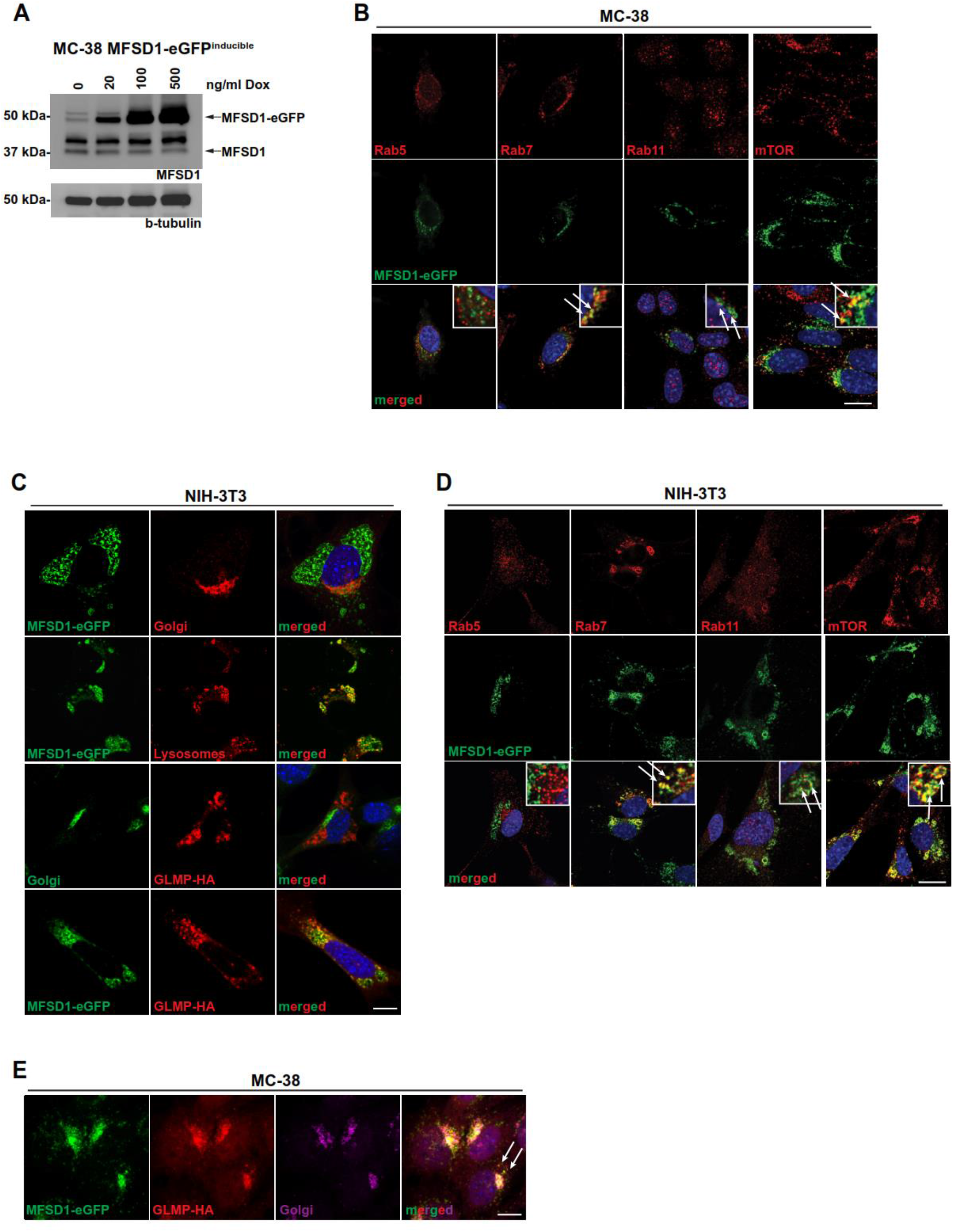
MFSD1-eGFP localizes to late-endosomes and lysosomes in MC-38 tumor cells and NIH-3T3 fibroblasts. **A)** MFSD1-eGFP expression was induced by different Doxycycline concentrations for 24 hours in MC-38 cells, and the expression of MFSD1-eGFP was analyzed by Western blotting. **B)** Immunofluorescence pictures of MC-38 MFSD1-eGFP cells with indicated markers. Co-localization (yes, partial, or no) is indicated at the bottom of each staining. Arrows indicate co-localization with indicated markers. **C)** Immunofluorescence pictures of NIH-3T3 MFSD1-eGFP cells with indicated markers. Arrows indicate co-localization with indicated markers. **D)** Immunofluorescence images of NIH-3T3 MFSD1-eGFP cells with indicated markers. **E)** Immunofluorescence pictures of MC-38 MFSD1-eGFP&GLMP-HA cells with Golgi-apparatus marker. Arrows indicate vesicular co-localization of MFSD1-eGFP and GLMP-HA. Bar = 10 μm. Arrows indicate co-localization.

**Supplementary Figure 3.**
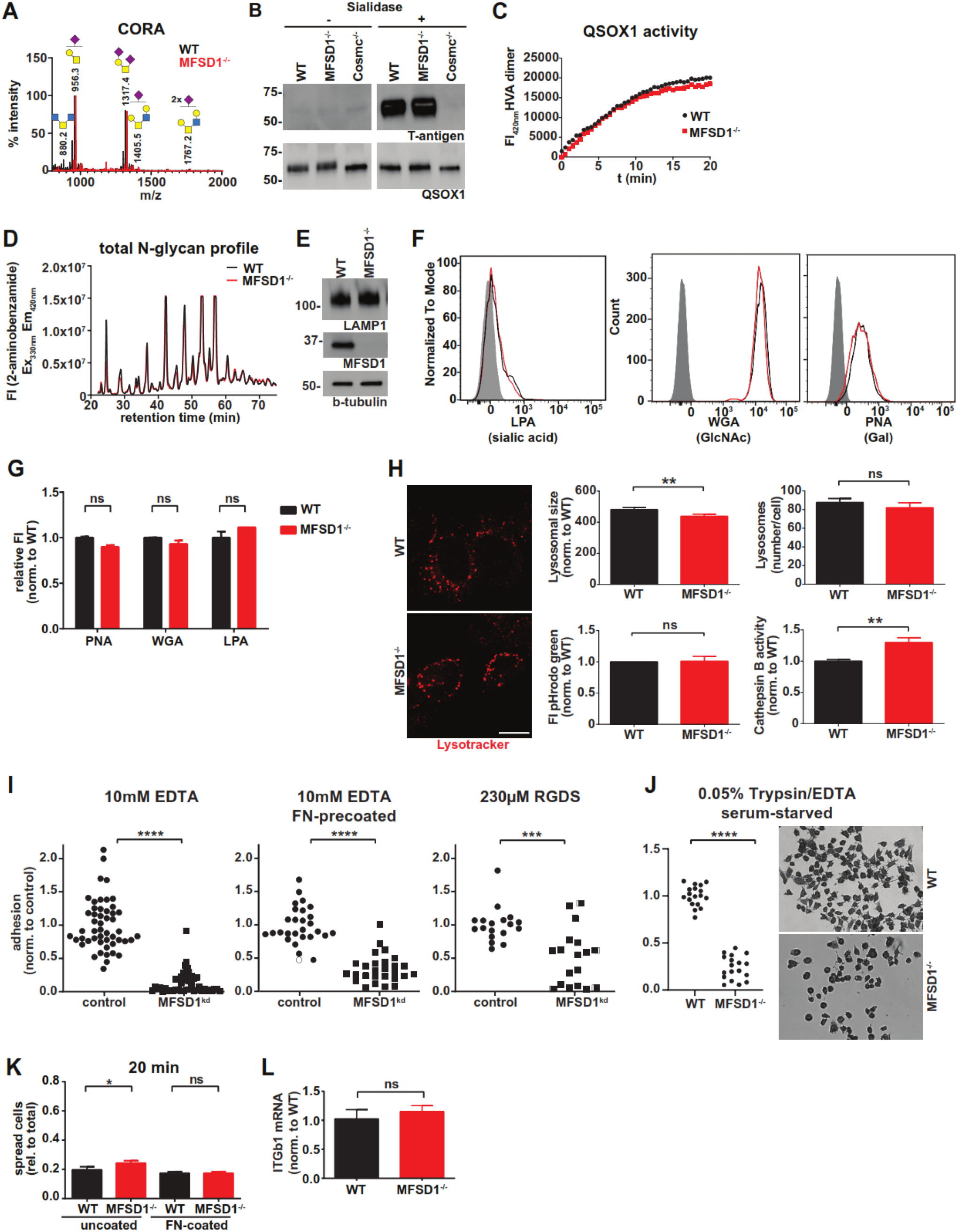
MFSD1 does not strongly affect glycosylation nor lysosomes, but rather the adhesive strength of MC-38 tumor cells. **A)** CORA (Cellular O-Glycome Reporter/Amplification) analysis by mass spectrometry of secreted O-glycans by MC-38 WT and MFSD1^-/-^ cells. The MFSD1^-/-^ diagram was shifted to the right to visualize the identical relative abundance of sialyl- and disialyl- T antigen in WT and MFSD1^-/-^ MC-38 cells. **B)** Western blot of purified QSOX1-StrepTagII purified from supernatant of MC-38 WT, MFSD1^-/-^, and Cosmc^-/-^ cells (n=2). **C)** QSOX1 activity measurement. The result of one out of two experiments is shown. **D)** LC profile of total N-glycans isolated from MC-38 WT and MFSD1^-/-^ cells. **E)** Western blot of LAMP1 in MC-38 WT and MFSD1^-/-^ cells. **F-G)** FACS analysis of cell surface lectin staining in MC-38 WT and MFSD1^-/-^ cells (n=2 for LPA; n=3 for PNA and WGA). **H)** LysoTracker staining of MC-38 WT and MFSD1^-/-^ cells. The listed parameters were analyzed by ImageJ. Lysosomal size (n>200), lysosomes number/cell (n>50), pHrodo green (n=4), Cathepsin B activity (n=6).Bar = 10 μm. n= **I-J)** MC-38 cell detachment with the indicated chemicals. Per experiment 3 view fields per replicate (three replicates per experiment) were analyzed. Adhesion was determined by the area covered by still adherent cells present on the cell culture plates, within experiments adhesion was normalized to that of the WT cells. **K)** Spreading assay of MC-38 WT and MFSD1^-/-^ on uncoated and FN-coated plates at 20 min time point (n≥21 view fields). **L)** qPCR analysis of β1-integrin transcription in MC-38 WT and MFSD1^-/-^ cells. GAPDH served as a control (n=3). * = p<0.05; ** = p<0.01; *** = p<0.001; **** = p<0.0001; ns = not significant.

**Supplementary Figure 4.**
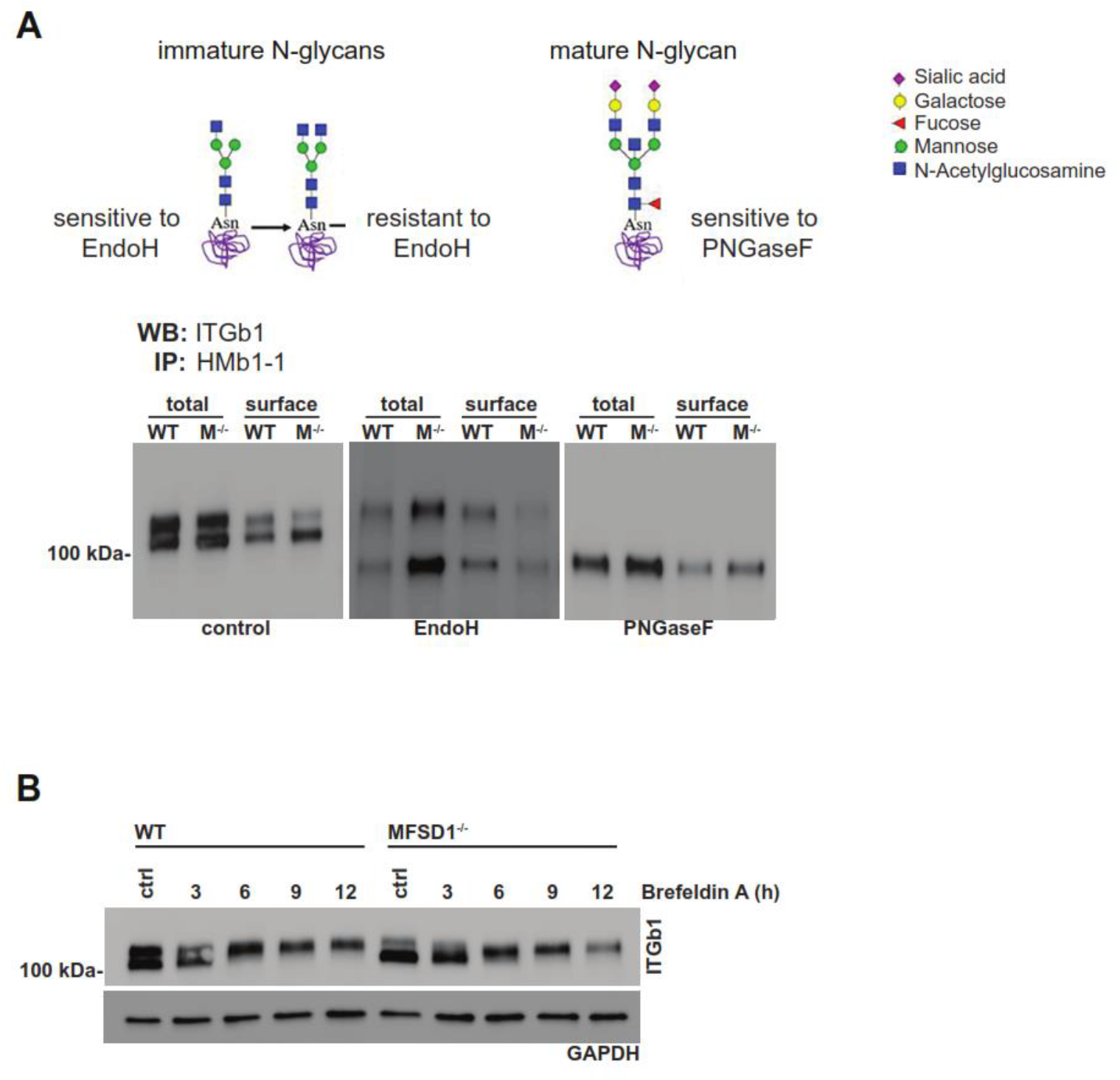
Immaturely N-glycosylated β1-integrin is sensitive to EndoH treatment. **A)** Schematic of N-glycans senstitive or resistant to EndoH or PNGaseF glycosidases (top panel). Western blot of immunoprecipitated β1-integrin from cell lysates (total) or the cell surface of MC-38 WT and MFSD1^-/-^ cells treated with glycosidase EndoH, PNGaseF, or untreated (control) (bottom panel) (n=1). **B)** Western blot of Brefeldin A treated MC-38 WT and MFSD1^-/-^ cell lysates (n=2).

**Supplementary Figure 5.**
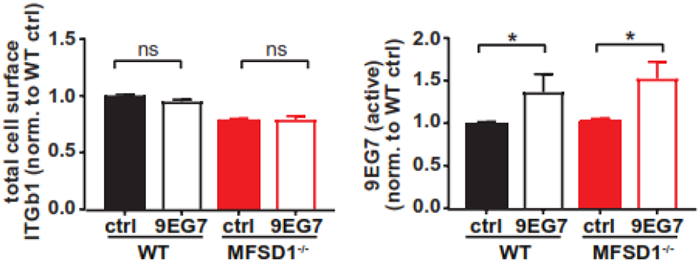
The increased β1-integrin activation index enables pro-metastatic phenotypes. **A)** Analysis of flow cytometry stainings (n=4).

